# Facilitation and inhibition of firing activity and N-methyl-D-aspartate-evoked responses of CA1 hippocampal pyramidal cells by alpha7 nicotinic acetylcholine receptor selective compounds in vivo

**DOI:** 10.1101/576520

**Authors:** Zsolt Kristóf Bali, Lili Veronika Nagy, Dénes Budai, István Hernádi

**Affiliations:** Department of Experimental Zoology and Neurobiology, Faculty of Sciences, University of Pécs, Pécs, Hungary, Postal address: 6 Ifjúság út, H-7624, Pécs, Hungary; János Szentágothai Research Center, Center for Neuroscience, University of Pécs, Pécs, Hungary, Postal address: 20 Ifjúság út, H-7624, Pécs, Hungary; Kation Scientific LLC, Minneapolis, Minnesota, United States of America, Postal address: 2010 East Hennepin Avenue Suite 8-103, Minneapolis, MN 55413, USA; Institute of Physiology, Medical School, University of Pécs, Pécs, Hungary, Postal address: 12 Szigeti út, H-7624, Pécs, Hungary; Grastyán Endre Translational Research Center, University of Pécs, Pécs, Hungary, Postal address: 6 Ifjúság út, H-7624, Pécs, Hungary

**Author notes:** Address for correspondence: University of Pécs, Department of Experimental Zoology and Neurobiology 6 Ifjúság út, H-7624, Pécs, Hungary.

**Keywords:** electrophysiology, acetylcholine, N-methyl-D-aspartate, nicotinic acetylcholine receptor, receptor desensitization, hippocampus

## Abstract

Alpha7 nicotinic acetylcholine receptors (nAChRs) are promising novel targets for the treatment of neurocognitive disorders. Although the cognitive enhancer potential of alpha7 nAChR agonists and positive allosteric modulators (PAMs) has been confirmed in several preclinical animal models, there are only sparse *in vivo* electrophysiological data on their effects on the firing activity and excitability of neurons. The present study investigated and compared local effects of alpha7 nAChR agonist PHA-543613 and PAMs PNU-120596 and NS-1738 on the spontaneous and N-methyl-D-aspartate-evoked (NMDA-evoked) firing rate of rat CA1 hippocampal pyramidal cells, *in vivo*. Furthermore, effects of alpha7 nAChR antagonist methyllycaconitine (MLA) and GABA were also tested. Results showed substantially different effects of the alpha7 nAChR agonist and PAMs. While PNU-120596 and NS-1738 predominantly and significantly increased both spontaneous and NMDA-evoked firing rate of the neurons, application of PHA-543613 resulted in almost equal distribution of facilitatory and inhibitory effects. The increase of the NMDA-evoked firing rate exerted by NS-1738 was superadditive over the sum of the single effects of NMDA and NS-1738. The simultaneous application of alpha7 nAChR agonist PHA-543613 and PAM NS-1738 resulted in additive increase of both spontaneous and NMDA-evoked firing rate. However, NS-1738 counteracted inhibitory effects of PHA-543613 in 5 out of 6 neurons, resulting in a synergistic potentiation of their firing responses to NMDA. Our results suggest that alpha7 nAChR PAMs increase neuronal excitability more potently than agonists, while the remarkable occurrence of inhibitory effects of PHA-543613 (possibly originating from receptor desensitization) implies that agonists may exert neuroprotective effects.

## 1. Introduction

In the central nervous system, the interplay of glutamatergic and cholinergic neurotransmission has been suggested to play a key role in various cognitive functions, especially in learning and memory. Modulation of cholinergic neurotransmission has also become the most widely used pharmacological strategy in the treatment of neurocognitive disorders, including Alzheimer’s disease (AD). For instance, cognitive symptoms of AD are improved by the inhibition of acetylcholinesterase (AChE) enzyme activity (McGleenon *et al.*, 1999) and by the modulation of N-methyl-D-aspartate glutamate receptor (NMDAR) signaling with the weak NMDAR antagonist memantine (Parsons *et al.*, 2007). Recently, direct stimulation of the alpha7 nicotinic acetylcholine receptor (nAChR) has also become a promising target to attenuate cognitive decline in dementia as they are highly expressed in brain areas essential for cognition, attention and working memory, including the hippocampus, cerebral cortex and limbic structures (Gotti *et al.*, 2006; Wallace and Porter, 2011). Alpha7 nAChRs are implicated in mediating fast synaptic transmission, neurotransmitter release and synaptic plasticity. In the hippocampus, alpha7 nAChRs are localized on interneurons and glutamatergic pyramidal cells in which they are widely expressed both presynaptically and postsynaptically (Fabian-Fine *et al.*, 2001; Dani and Bertrand, 2007).

Recently, a great number of agonists, antagonists and positive allosteric modulators (PAMs) of the alpha7 nAChRs have been designed with considerable emphasis on developing approved cognitive enhancer substances for clinical use. In general, while agonists directly activate the nAChR, PAM compounds rather increase the potency of endogenous nicotinic agonists, such as ACh (Wallace and Porter, 2011), through modulating the positive allosteric site of the receptor. Hence, the use of PAMs may offer an alternative way to activate alpha7 nAChRs.

The activation of alpha7 nAChRs by agonists or PAMs has already been shown to be effective to enhance learning and memory in several behavioral tests in preclinical animal models of various cognitive disorders (Ng *et al.*, 2007; Timmermann *et al.*, 2007; Bali *et al.*, 2015; Nikiforuk *et al.*, 2015; for a review see Wallace and Porter, 2011). Moreover, both alpha7 nAChR agonists and PAMs have been reported to show interaction with the effects of memantine in behavioral experiments (Nikiforuk, Potasiewicz, *et al.*, 2016; Bali *et al.*, 2019). However, the effect of alpha7 nAChR agonists and PAMs on the electrophysiological properties of neurons has not yet been fully characterized and compared in light of their cognitive enhancer potency. Although electrophysiological properties of alpha7 nAChRs have been extensively studied *in vitro* (Hurst *et al.*, 2005; Dinklo *et al.*, 2011; Maex *et al.*, 2014), little is known about their *in vivo* response to direct stimulation.

A recent electrophysiological study from our laboratory has demonstrated the role of alpha7 nAChRs in the synergistic action of ACh and NMDA in the hippocampal CA1 area *in vivo* (Bali *et al.*, 2017). Namely, we found that the synergistic effect of ACh and NMDA on the firing activity of the neurons was robustly decreased by the systemic administration of alpha7 nAChR antagonist methyllycaconitine (MLA) but not by muscarinic antagonist scopolamine. In summary, this prior work provided an *in vivo* evidence that alpha7 nAChRs may contribute to pyramidal cell activation through the potentiation of glutamatergic neurotransmission.

In the present study, our aim was to examine whether exogenous alpha7 nAChR agonists and PAMs exert the same or similar effects as ACh on the firing activity of hippocampal CA1 neurons, and whether and how exogenous alpha7 nAChR ligands may also potentiate firing rate responses to glutamatergic receptor stimulation. We also aimed to compare the effect and efficacy of different alpha7 nAChR ligands and examine possible agonist-PAM interactions under *in vivo* conditions in the hippocampus, a brain structure highly relevant to declarative memory formation and memory consolidation. Therefore, extracellular firing activity of rat hippocampal CA1 neurons was recorded, and the effects of different locally applied alpha7 nAChR ligands (agonist PHA-543613, PAMs PNU-120596 and NS-1738, and antagonist MLA) were tested on the spontaneous and NMDA-evoked firing activity of the neurons. The present experiments provide new insights into the actions of alpha7 nAChR ligands on the neuronal level *in vivo*, and into the differences between the modulation of alpha7 nAChR by agonists and PAMs with implications on their potential therapeutic use in the future.

## 2. Methods

### 2.1. Animals and surgery

This study was approved by the Animal Care Committee of the University of Pécs (licence no. BA02/2000-80/2017). Procedures fully complied with Decree No. 40/2013 (II. 14.) of the Hungarian Government and EU Directive 2010/63/EU on the protection of animals used for scientific purposes. Altogether 46 adult Wistar rats (31 male, 15 female, Charles River Laboratories, Germany) were used in the experiments. The animals were housed in a conventional animal care facility under a 12:12 h light/dark cycle under controlled conditions (22±2 °C temperature, and 40 to 60% humidity). Food and water were available *ad libitum*. Surgical preparations were carried out as described previously (Bali *et al.*, 2014, 2017). Rats were initially anesthetized using chloral hydrate (400 mg/kg b.w., i.p.), and stable anesthesia was further maintained by continuous intravenous administration of the anesthetic via a jugular vein cannula (chloral hydrate, 100 mg/kg/h initially, later adjusted if needed). Anesthesia was continuously maintained until the end of recordings, then, animals were humanely euthanized by an intravenous overdose of pentobarbital (Euthanimal 40%, Alfasan, Woerden, Netherlands). Surgery was performed in a stereotaxic frame. First, an incision was made on the scalp, then a hole was drilled in the skull. A small part of the dura mater was removed, and a multi-barrel carbon fiber microelectrode (Carbostar, Kation Scientific Ltd., Minneapolis, MN, USA) was inserted into the CA1 area of the dorsal hippocampus according to the rat brain atlas by Paxinos and Watson (2014). The relevant coordinates were as follows: AP 3.5-5.5 from bregma, ML 1.5-2.8 from sagittal suture, and DV 2.0-4.3 from dura. The microelectrode consisted of a carbon fiber recording channel (∼7 µm in diameter, with a 25 µm long active electrode tip), and 3 to 6 glass capillaries (∼1 µm in tip diameter each) surrounding the central channel for microiontophoretic application of different compounds. The microiontophoretic capillaries ended at the same level as the borosilicate glass insulation of the carbon fiber (Budai and Molnár, 2001).

### 2.2. Extracellular spike recording and microiontophoresis

Electrophysiological signal was recorded extracellularly through the central carbon fiber of the microelectrode. The signal was amplified and band-pass filtered between 300 Hz and 3000 Hz using analog electrophysiological amplifiers (BioAmp, Supertech Ltd., Pécs, Hungary; NeuroLog, Digitimer Ltd., Welwyn Garden City, UK). The signal was digitized at 25 kHz using CED Power 1401 analog-to-digital converter and Spike2 software (Cambridge Electronic Design Ltd., Cambridge, UK). The micropipettes of the microelectrodes were filled with some of the solutions of the following pharmacological compounds (manufacturer, pipette concentrations and microiontophoretic ejection currents are in parentheses): NMDA (Sigma-Aldrich, 50mM, −10 to −75 nA), PHA-543613 hydrochloride (Tocris, 50 mM, +20 to +200 nA), PNU-120596 (Tocris, 40 mM, −10 to −100 nA), NS-1738 (Tocris, 40 mM, −10 to −130 nA), MLA (Sigma-Aldrich, 20 mM, +20 to +80 nA), GABA (Tocris, 20 mM, +20 to +80 nA). The amount of the compound delivered with microiontophoresis was proportional to the applied current and to the length of ejection. During recordings, pharmacological compounds were locally delivered from the micropipettes in the vicinity of the recorded neurons by means of microiontophoresis. For microiontophoretic delivery, a constant current generator was used (Neurophore BH-2 System, Medical System Corp., Greenvale, NY, USA). Between drug administrations, a low retention current with opposite charge was applied on the microiontophoresis pipettes to avoid leakage of compounds.

Extracellular action potentials (spikes) were extracted offline from the recordings using the Spike2 software. Spikes were defined as sharp (∼1 ms in width) electrophysiological signals with an amplitude at least 5 times larger than the root-mean-square of the background noise. Then, spikes were further sorted in single-spiking and complex-spiking neuronal subtypes in the Klusters software (Lynn Hazan, Buzsáki lab, Rutgers, Newark, NJ, USA, RRID:SCR_008020) according to their shape and the auto-correlograms representing the firing pattern of the neurons. Conversion of data between Spike2 and Klusters was performed using our custom made scripts freely available on Mendeley Data (Bali *et al.*, 2014; Bali, 2018a; b). At every discharge event, complex-spiking neurons typically fired between 2 to 7 spikes in a train with short inter-spike intervals (3 to 6 ms), while single-spiking neurons fired with longer inter-spike intervals (>6 ms). Therefore, spike clusters that showed sharp peaks between 3 and 6 ms on the autocorrelograms were considered complex-spiking neurons. According to Csicsvari et al. (1998), complex-spiking neurons were considered as pyramidal cells of the CA1 area. For detailed description of the separation process, refer to Bali *et al.* (2014). In this study, only complex-spiking neurons were analyzed, spike clusters that did not show complex-spiking behavior were excluded from further analysis.

### 2.3. Experimental protocol

Throughout the recording sessions, NMDA was periodically ejected for 5 s in every 120 s. Ejection currents for NMDA was set to evoke a significant increase of the neuronal firing rate during NMDA application (NMDA-peaks). After recording three consecutive 120 s long periods with stable NMDA-peaks, one of the test compounds (PHA-543613, NS-1738, PNU-120596, MLA or GABA) was ejected microiontophoretically for 70 s (starting 60 s before the next NMDA-peak). In order to investigate the interactions between the alpha7 nAChR agonist PHA-543613 and the PAM NS-1738, the effect of the two compounds was also assessed during their simultaneous application. The experimental protocol and the time intervals for the calculation of firing rate values are represented on Fig. 1A. Firing rate values were determined for each pharmacological trial (i.e., microiontophoretic application of a test compound). Control spontaneous firing rate (Sp) was calculated in the period before the microiontophoretic application of a test compound and was defined as the firing rate (1/s, Hz) measured in a 60 s time window immediately before the next NMDA-peak. Control NMDA-evoked firing rate (NMDA) was defined by averaging the NMDA-evoked firing rate during the three NMDA-peaks preceding the application of the test compound. The effect of a test compound on the spontaneous firing rate of the neurons (Sp_Drug) was assessed by calculating the firing rate during the application of the compound in a 60 s time window immediately before the next NMDA-peak. The effect of the test compound on the NMDA-evoked firing rate (NMDA_Drug) was assessed by calculating the firing rate during the simultaneous application of NMDA and the test compound. Effects of test compounds on spontaneous and NMDA-evoked firing were also expressed in the percentage of the corresponding control value (further referred to as normalized spontaneous and NMDA-evoked firing rates, calculated as: (Sp_Drug/Sp)×100; and (NMDA_Drug/NMDA)×100, respectively).

**Figure 1.**
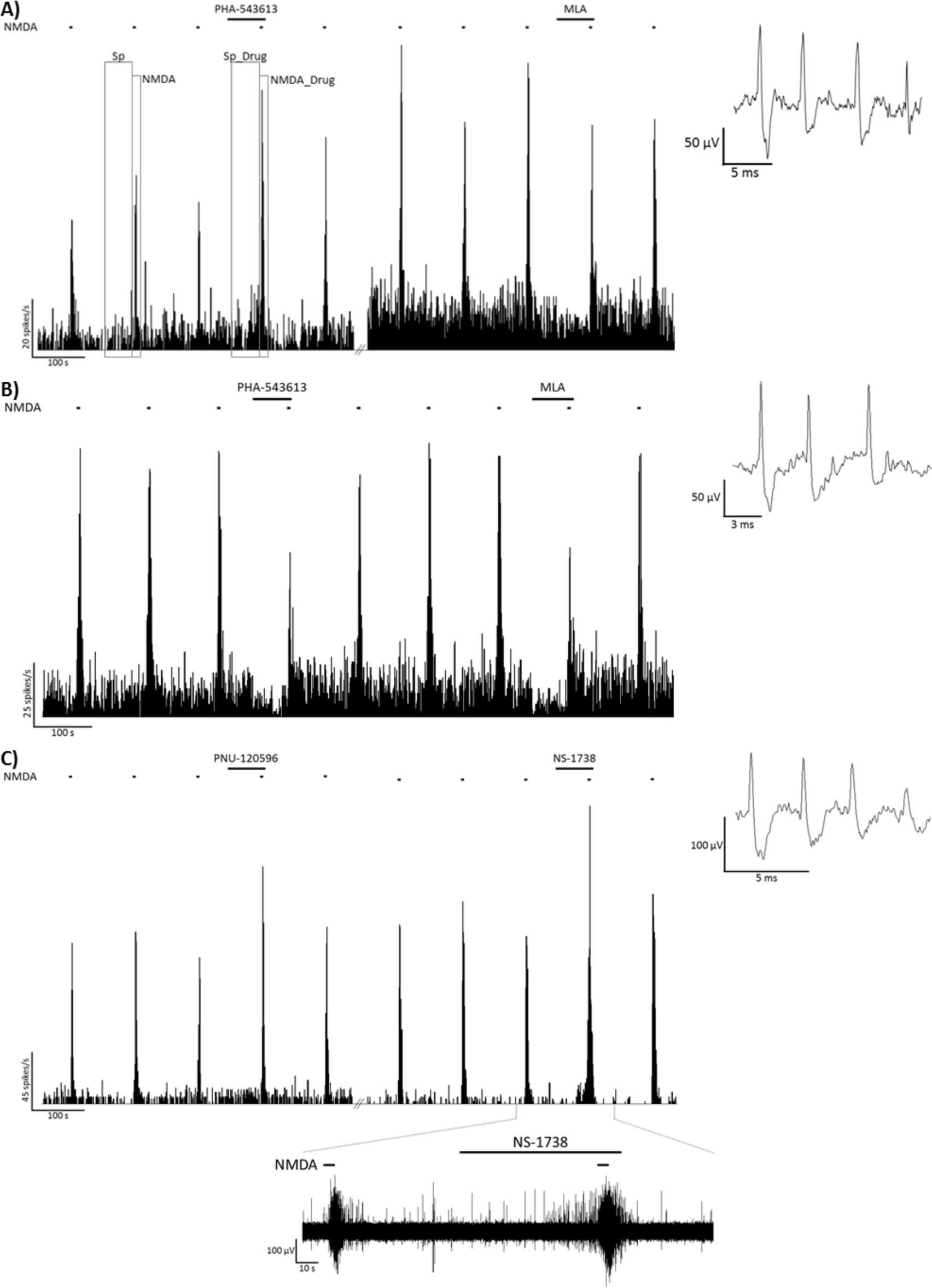
Firing rate histograms from representative recordings of the local effects of PHA-543613 and MLA (A, B), and PNU-120596 and NS-1738 (C) on the spontaneous and NMDA-evoked firing activity of CA1 hippocampal pyramidal neurons. Furthermore, panel A shows the time windows for calculating spontaneous and NMDA-evoked firing rates before (Sp, NMDA, respectively), and during (Sp_Drug, NMDA_Drug) microiontophoretic application of a given test compound. Insets to the right show a typical extracellular action potential recorded in the given experimental session. Bottom inset on panel C shows the raw electrophysiological recording in the time window marked on the horizontal axis of the corresponding histogram.

### 2.4. Statistical analysis

In the statistical analysis, neurons were chosen as units of observation. Therefore, we determined a single value for every neuron for each measured parameter from the data obtained in individual trials. Namely, firing rate values in control states (Sp, NMDA) and during the application of test compounds (Sp_Drug, NMDA_Drug) were averaged between trials. Normalized (percentage) firing rate as an effect of the test compound on the neuron was defined as the median of data in individual trials (in order to avoid the bias caused by outlier values). Furthermore, on the basis of the median normalized firing rate, neurons were sorted in groups showing the increase, decrease or no change of spontaneous or NMDA-evoked firing rate as a result of the application of the given test compound. Namely, a response of a neuron to a given test compound was considered as firing rate increase if the median normalized firing rate was above 120%, while the response of the neuron was considered as firing rate decrease if the median normalized firing rate was below 80%. A normalized firing rate between 80% and 120% was considered as no effect.

The overall effect of a given test compound was determined by comparing the number of neurons showing firing rate increase and decrease using two-tailed binomial test (neurons that did not respond to the treatments were excluded from this analysis). If the binomial test was significant (p<0.05), the effect (i.e., firing rate increase or decrease) occurring with higher incidence was considered as the general effect of the compound. A non-significant binomial test showed the equal distribution of firing rate increasing and decreasing effects among the neuronal population. The effect of a given compound was also assessed using the normalized firing rate values by comparing them to the baseline value (100%) using one-sample Wilcoxon signed rank test. Furthermore, the effects of different compounds were also compared by testing the differences between normalized firing rate data after different treatments using the Kruskal-Wallis test and post-hoc Mann-Whitney U test corrected with Holm’s method for multiple comparisons (Holm, 1979). In experiments testing the effect of simultaneous application of PHA-543613 and NS-1738, the effects of the compounds were assessed by comparing the control firing rate with the firing rate measured during the application of one or both compounds using the Wilcoxon signed rank test for paired samples.

We further analyzed the nature of interactions between different test compounds. First, we tested whether the effect of a given test compound on the NMDA-evoked firing rate is significantly higher than the sum of the effects of NMDA and the test compound alone on the spontaneous firing rate. A similar analysis was described in our previous study to test additive/superadditive interaction between the effect of NMDA and ACh (Bali *et al.*, 2017). Briefly, we calculated the sum of the individual firing rate changing effects of NMDA and the given test compound relative to the spontaneous firing rate using the following equation: (Sp_Drug-Sp)+(NMDA-Sp). Then, we calculated the relative firing rate change resulted from the simultaneous application of NMDA and the test compound (combined effect): NMDA_Drug-Sp. Similarly, in experiments investigating the interactions between PHA-543613 and NS-1738, we compared the sum of the individual effects of PHA-543613 and NS-1738 to their combined effect during simultaneous application. Thus, the following comparison was made to assess interactive effects on the spontaneous firing rate:

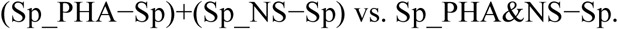

A similar comparison was made to assess the interactive effects on the NMDA-evoked firing rate:

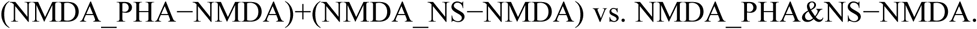

In both interaction analyses, the sum of the individual effects were compared with the combined effect using the Wilcoxon signed rank test for paired samples. A significantly higher relative change during the simultaneous application of the two compounds suggested a superadditive (synergistic) interaction between the two compounds in increasing the firing rate of neurons, while a combined effect significantly lower than the sum of individual effects suggested an antagonistic interaction. A non-significant result implies a potential additive relationship between the effects of the two test compounds.

## 3. Results

The effects of local iontophoretic application of alpha7 nAChR agonist PHA-543613, PAMs NS-1738 and PNU-120596, and antagonist MLA were tested on the spontaneous and NMDA-evoked firing activity of altogether 111 hippocampal pyramidal neurons. Furthermore, the effects of GABA were tested as a control on 19 neurons (from 8 rats; data not shown on figures). Figure 1 shows examples of the pharmacological effects of alpha7 nAChR ligands on representative recordings. Figure 2 summarizes the distribution of the number of neurons that increased, decreased or did not change their firing rate during the application of the given compound. Furthermore, the comparison of normalized firing rates during different treatments with alpha7 nAChR ligands is shown on Fig. 3.

**Figure 2.**
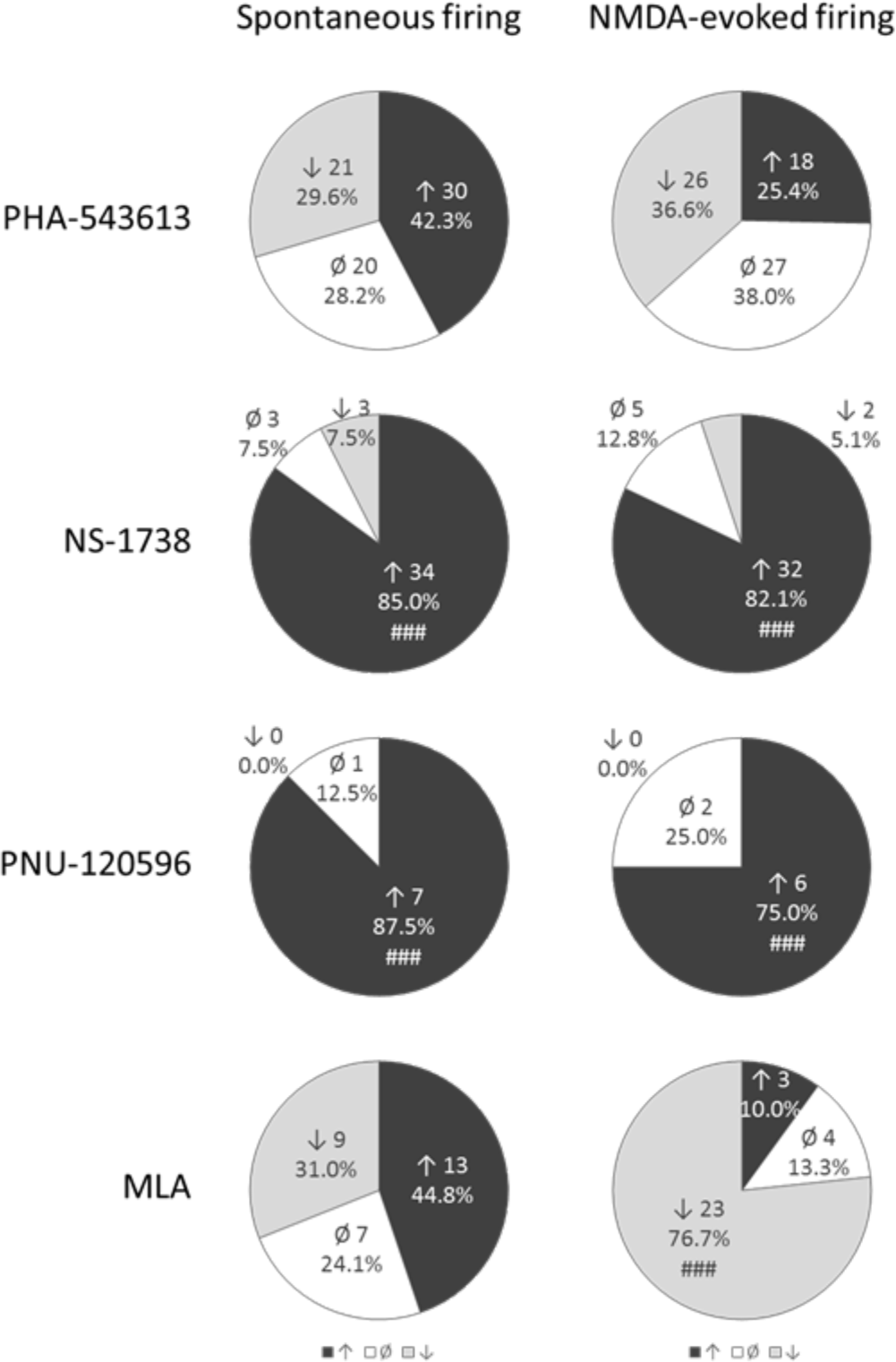
Distribution of the number of neurons responded with firing rate increase, decrease or no response to the microiontophoretic application of the given test compound. Hash symbols indicate effects occured with significantly higher probability then the opposite effect (i.e., firing rate increase or decrease) according to binomial test: ### p<0.001.

**Figure 3.**
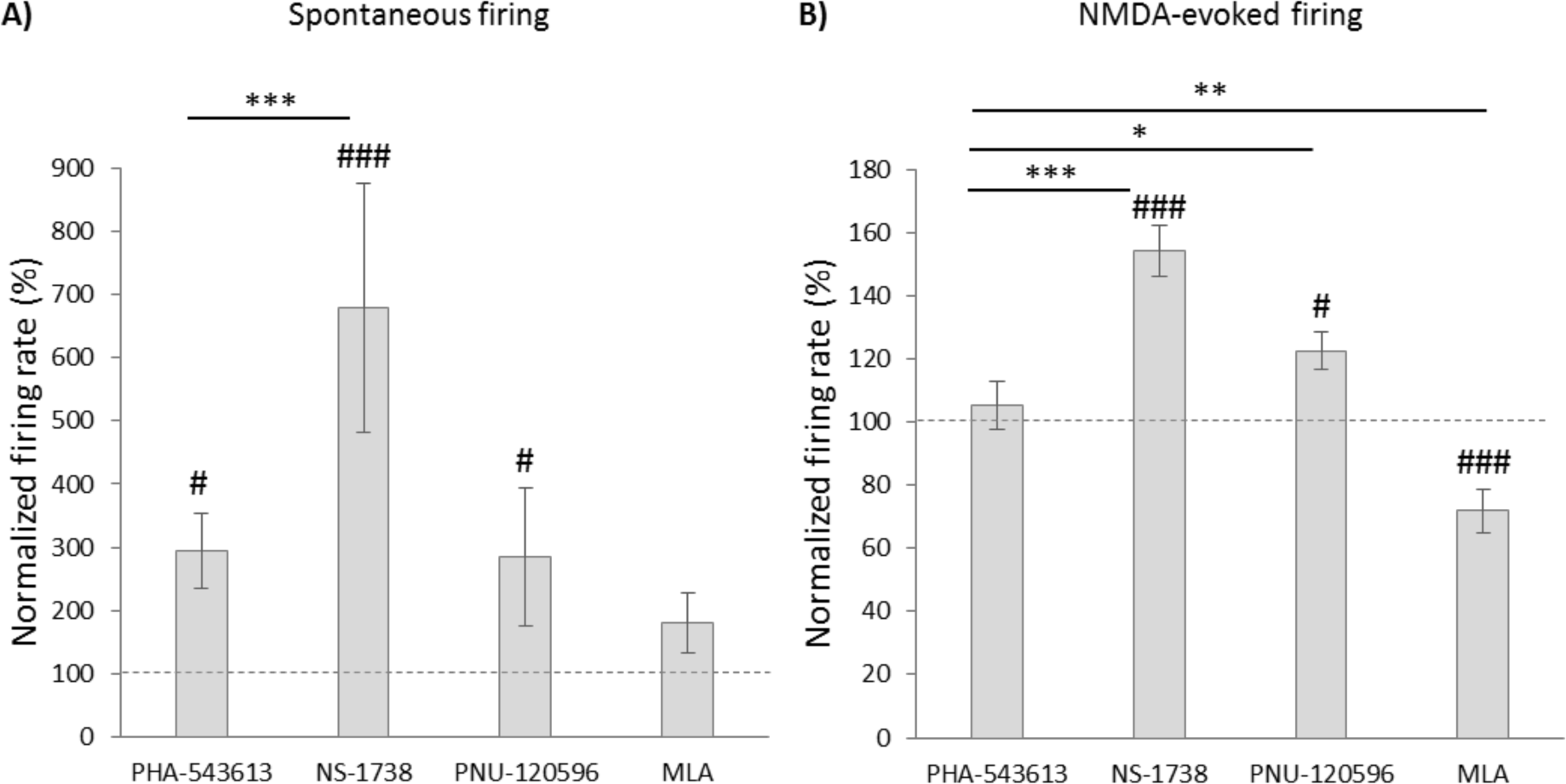
Comparison of the normalized spontaneous (A) and NMDA-evoked (B) firing rate during the local application of different test compounds (mean±SEM). Effects of PHA-543613, NS-1738, PNU-120596 and MLA on the spontaneous firing rate were tested on 71, 40, 8 and 29 neurons, respectively (Kruskal-Wallis H=21.1, p<0.001). Effects of PHA-543613, NS-1738, PNU-120596 and MLA on the NMDA-evoked firing rate were tested on 71, 39, 8 and 30 neurons, respectively (Kruskal-Wallis H=49.3, p<0.001). Hash symbols above bars indicate that the given treatment resulted in a normalized firing rate significantly higher than the baseline (100%): # p<0.05, ### p<0.001. Asterisks indicate significant differences between the mean normalized firing rates as an effect of different test compounds (post-hoc comparisons): * p<0.05, ** p<0.01, *** p<0.001.

### 3.1. Effects of test compounds on spontaneous firing activity

The mean spontaneous firing activity of the recorded neurons was 2.92±0.37 Hz (mean±SEM) in baseline conditions. Baseline firing rate of neurons was not different in male and female rats (2.69±0.41 vs. 3.66±0.84, respectively; Mann-Whitney U test: U=1218.5, z=1.045, p=0.296).

Both firing rate increasing and decreasing effects were observed on a remarkable number of neurons during the iontophoresis of alpha7 nAChR agonist PHA-543613. Representative recordings of firing rate increasing and decreasing effects of PHA-543613 are shown on Fig. 1A and Fig. 1B, respectively. In 30 neurons out of 71 (42.3%) PHA-543613 increased the spontaneous firing rate, while 21/71 neurons (29.6%) were found that responded with firing rate decrease to PHA-543613 application (increasing vs. decreasing effects: 30 vs. 21, binomial test: p=0.262; Fig. 2). Thus, the distribution of neurons responding with firing rate increase and decrease was almost equal. However, the normalized firing rate was significantly higher during PHA-543613 application compared with the baseline (293.3±59.2% vs. 100%, W=1635.5, z=2.048, p=0.041).

On the contrary, alpha7 nAChR PAMs predominantly increased the spontaneous firing rate of neurons. The great majority of the neurons responded with the increase of the firing rate to the application of both NS-1738 and PNU-120596 (34/40, 85%, and 7/8, 87.5%, respectively), resulting in a significantly higher prevalence of firing rate increasing effects compared to firing rate decreasing effects (NS-1738: 34 vs. 3 neurons, binomial test: p<0.001; PNU-120596: 7 vs. 0 neurons, p<0.001; Fig. 2). The spontaneous firing rate of the neurons significantly increased during the application of both PAM compounds compared with the baseline (NS-1738: 679.6±197.8%, W=784, z=5.027, p<0.001; PNU-120596: 284.6±110.1%, W=36, z=2.521, p=0.012). The normalized firing rate during the application of NS-1738 was significantly higher than that resulted from the application of PHA-543613 (Fig. 3A; Kruskal-Wallis H=21.1, p<0.001; NS-1738 vs. PHA-543613: p<0.001).

Interestingly, the effects of MLA were also distributed almost equally between firing rate increasing (13/29 neurons, 44.8%) and decreasing (9/29 neurons, 31.0%) effects, similar to PHA-543613 (increases vs. decreases: 13 vs. 9, binomial test: p=0.524; Fig. 2). On the other hand, the normalized firing rate was not significantly different from the baseline during MLA application (179.8±47.3%, W=277, z=1.685, p=0.092).

As a reference inhibitory compound, GABA decreased the spontaneous firing rate of all 18 examined neurons (increases vs. decreases: 0 vs. 18, p<0.001). The spontaneous firing rate decreased to 20.9±3.7% of the baseline activity as a result of GABA application (W=171, z=3.724, p<0.001).

### 3.2. Effects of test compounds on NMDA-evoked firing

The mean control NMDA-evoked firing activity of the recorded neurons was 29.04±1.56 Hz. Neurons recorded from males and females did not show difference in their responsiveness to NMDA (29.35±1.94 vs. 28.33±2.48, respectively; U=1410, z=0.2623, p=0.793).

Similar to the changes in spontaneous firing rate, PHA-543613 exerted both increasing and decreasing effects on the NMDA-evoked firing rate of neurons with almost equal distribution (increases: 18/71 neurons, 25.4%; decreases: 26/71 neurons, 36.6%; binomial test: 18 vs. 26, p=0.291; Fig. 2). Furthermore, the normalized firing rate during the application of PHA-543613 was not higher than the control baseline (105.5±7.6%, W=1385, z=0.834, p=0.404). Representative recordings of firing rate increasing and decreasing effects of PHA-543613 are shown on Fig. 1A and Fig. 1B, respectively.

On the contrary, PAM compounds predominantly increased the NMDA-evoked firing rate of the neurons (NS-1738: 32/39 neurons, 82.1%, increases vs. decreases: 32 vs. 2, p<0.001; PNU-120596: 6/8 neurons, 75.0%, increases vs. decreases: 6 vs. 0, p<0.001; Fig. 2) to a normalized firing rate significantly higher than the control baseline (NS-1738: 154.5±8.1%, W=750, z=5.024, p<0.001; PNU-120596: 122.6±5.8%, W=34, z=2.240, p=0.025). Application of PAMs resulted in significantly higher normalized NMDA-evoked firing rate than PHA-543613 (Fig. 3B; Kruskal-Wallis H=49.3, p<0.001; NS-1738 vs PHA-543613: p<0.001; PNU-120596 vs. PHA-543613: p=0.049). Representative recordings of the firing rate increasing effects of NS-1738 and PNU-120596 are shown on Fig. 1C.

Methyllycaconitine exerted a robust inhibitory effect on the NMDA-evoked firing rate by decreasing it in 23 out of 30 neurons (76.7%, increases vs. decreases: 3 vs. 23, p<0.001; Fig. 2) to 72.0±6.9% of the control in average (W=402.5, z=3.500, p<0.001). Compared with PHA-543613, normalized NMDA-evoked firing rate during MLA application was significantly lower than during PHA-543613 application (MLA vs. PHA-543613: p=0.001).

Gamma-aminobutyric acid decreased the NMDA-evoked firing rate in 17 out of 19 neurons without any instances for firing rate increasing effects (increases vs. decreases: 0 vs. 17, p<0.001). The NMDA-evoked firing rate decreased to 38.1±7.1% of the control level (W=188, z=3.743, p<0.001) as a result of the application of GABA.

### 3.3. Assessment of interactions between NMDA and test compounds

We analyzed the interaction of the effects of drugs on the spontaneous firing rate and on the NMDA-evoked firing rate (Fig. 4). We compared the sum of the effects of NMDA and different drugs with the effect of the simultaneous application of NMDA and the given drug on the firing rate ((NMDA-Sp)+(Sp_Drug-Sp) vs. NMDA_Drug-Sp, respectively). The firing rate increase after the simultaneous application of NMDA and PHA-543613 (19.37±1.58 Hz) did not differ from the sum of the effects of separately iontophoretized NMDA and PHA-543613 (21.56±1.55 Hz, W=1607, z=1.885, p=0.059), implying that simultaneously applied NMDA and PHA-543613 exerted an additive effect on the firing rate. On the other hand, NS-1738 increased the firing rate in a superadditive manner when applied together with NMDA in comparison with the sum of their effects in single applications (NMDA_Drug-Sp vs. (NMDA-Sp)+(Sp_Drug-Sp): 27.09±2.27 Hz vs. 21.21±1.97 Hz, W=698, z=4.298, p<0.001). Moreover, the antagonistic interaction between NMDA and MLA was further confirmed by finding a significantly lower combined effect compared to the sum of the individual effects of NMDA and MLA (NMDA_Drug-Sp vs. (NMDA-Sp)+(Sp_Drug-Sp): 15.47±2.43 Hz vs. 25.80±2.81 Hz, W=408, z=4.120, p<0.001).

**Figure 4.**
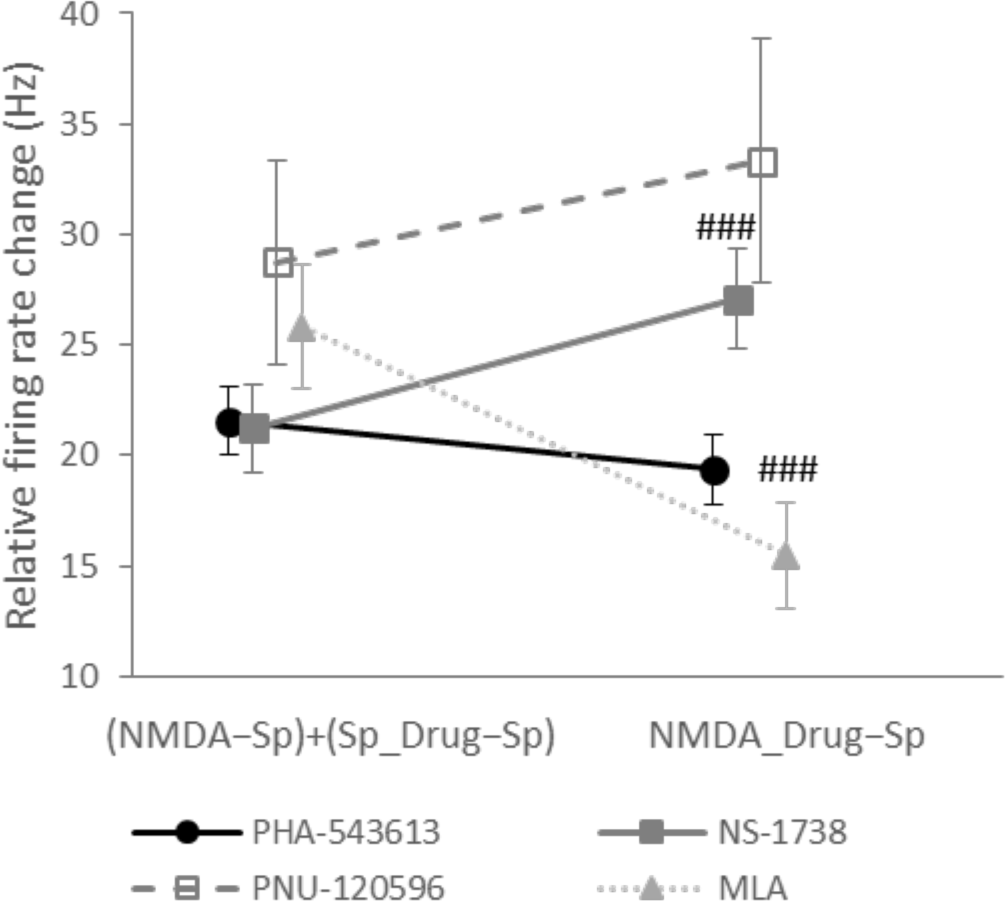
Assessment of the type of interaction between NMDA and different test compounds. Data points and error bars represent mean and SEM, respectively. Sample sizes were 71, 39, 8 and 29 for PHA-543613, NS-1738, PNU-120596 and MLA, respectively. Hash symbols indicate significant difference between the summarized effect of NMDA and the given test compound during their sole administration ((NMDA-Sp)+(Sp_Drug-Sp)) and their combined effect during their co-application (NMDA_Drug-Sp) according to Wilcoxon signed rank test: ### p<0.001.

### 3.4. Interactions between PHA-543613 and NS-1738

We tested the interactions between PHA-543613 and NS-1738 on 19 neurons by investigating their effects on spontaneous and NMDA-evoked firing rate after their single and combined application. Representative recordings and data plots on the effect of PHA-543613, NS-1738 and their simultaneous application are shown on Fig. 5. Figure 5A shows an example of simultaneous applications when PHA-543613 alone increased the spontaneous and NMDA-evoked firing rate, while Fig. 5B shows an example, where single application of PHA-543613 exerted an inhibitory effect.

**Figure 5.**
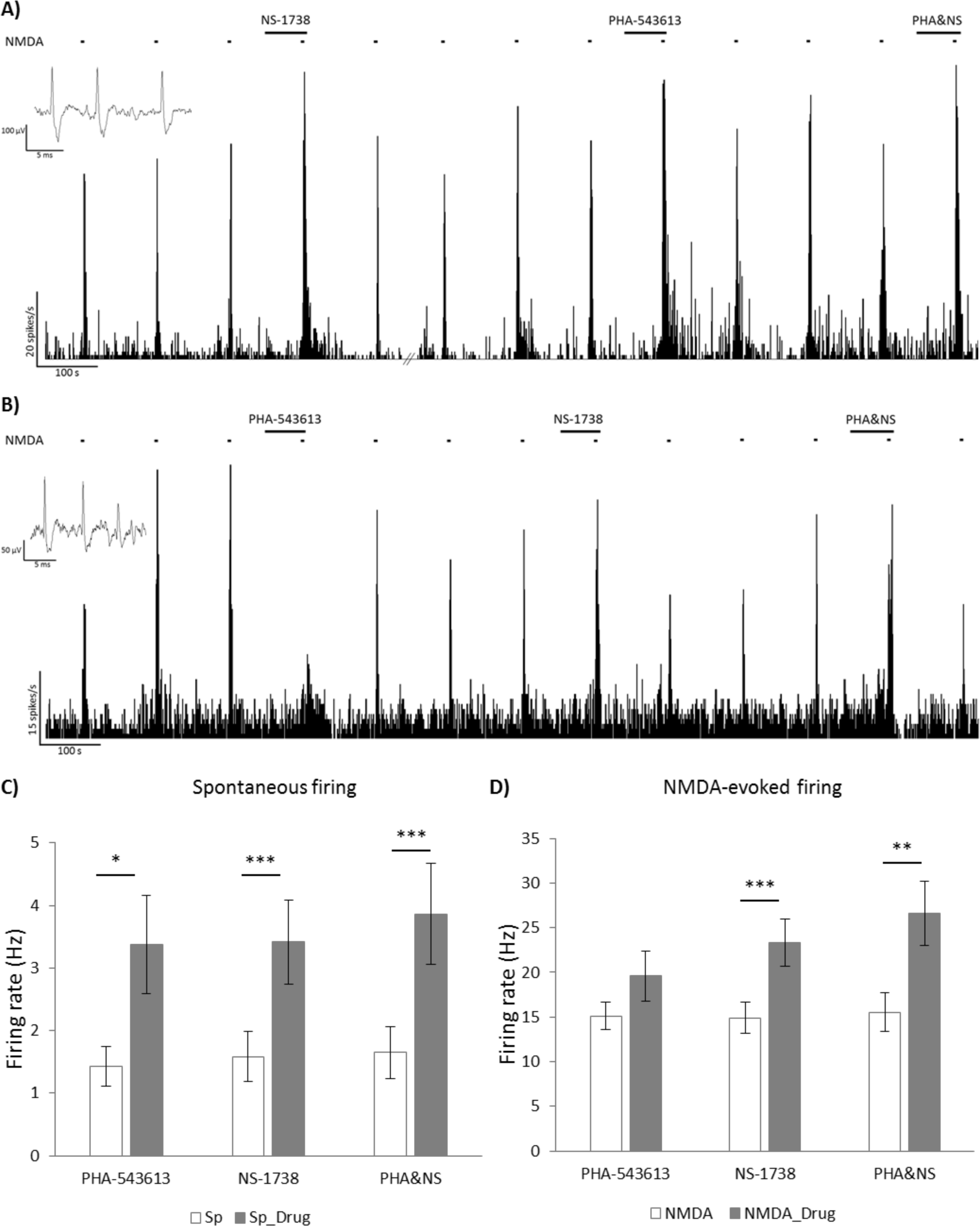
Comparison of the effects of alpha7 nAChR agonist PHA-543613 and PAM NS-1738 during their separate and simultaneous application. Firing rate histograms show typical examples of neurons that responded with firing rate increase (A) or with firing rate decrease (B) to the microiontophoretic application of PHA-543613. Insets show typical extracellular action potentials of CA1 hippocampal pyramidal cells recorded in the given experimental session. Bar charts (mean±SEM) assess the effects of PHA-543613, NS-1738 and their co-application on the spontaneous (C, N=20) and NMDA-evoked (D, N=19) firing rate. Asterisks indicate significant effects of the given compound compared with the baseline firing rate value according to Wilcoxon signed rank test: * p<0.05, ** p<0.01, *** p<0.001.

Out of the 19 neurons, single application of PHA-543613 increased the spontaneous firing rate of 12 neurons, while only 3 neurons showed decreased firing rate after PHA-543613 application. The spontaneous firing rate significantly increased from 1.4±0.3 Hz to 3.4±0.8 Hz (W=141, z=2.417, p=0.016, Fig. 5C). NS-1738 in single application increased the spontaneous firing rate in 17 out of 19 neurons (with firing rate decrease in 1 neuron), where the firing rate increased from 1.6±0.4 Hz to 3.4±0.7 Hz in average (W=198.5, z=3.492, p<0.001). The simultaneous application of PHA-543613 and NS-1738 resulted in the increase of spontaneous firing rate in 18 out of 19 neurons, and no decreasing effect was observed. The simultaneous application of PHA-543613 and NS-1738 also increased the firing rate of two neurons that responded with decreased firing activity to PHA-543613 alone. The combination of PHA-543613 and NS-1738 increased the spontaneous firing rate from 1.7±0.4 Hz to 3.9±0.8 Hz (W=207, z=3.808, p<0.001). The simultaneous application of PHA-543613 and NS-1738 did not resulted in a significantly higher relative increase of the spontaneous firing rate (combined effect) compared with the sum of the individual firing rate increasing effect of the two compounds (Sp_PHA&NS-Sp vs. (Sp_PHA-Sp)+(Sp_NS-Sp): 2.2±0.6 Hz vs. 3.8±0.9 Hz, W=143, z=1.419, p=0.156).

Application of PHA-543613 increased the NMDA-evoked firing rate in 9 out of 19 neurons, when applied alone, while the decrease of the NMDA-evoked firing rate was observed on 6 neurons. The NMDA-evoked firing rate after the single application of PHA-543613 (19.6±2.8 Hz) was not significantly higher than the control NMDA-evoked firing rate (15.1±1.5 Hz, W=127, z=1.288, p=0.198, Fig. 5D). NS-1738 increased the NMDA-evoked firing rate in 15 out of 19 neurons, and there was no neuron that responded with decreased NMDA-evoked firing rate to the single application of NS-1738. In average, NS-1738 significantly increased the firing rate of the neurons from 14.9±1.7 Hz to 23.3±2.7 Hz (W=190, z=3.823, p<0.001). The simultaneous application of PHA-543613 and NS-1738 increased the NMDA-evoked firing rate in 16 out of 19 neurons, while 2 neurons responded with decreased NMDA-evoked firing rate to the combined application. The NMDA-evoked firing rate significantly increased from 15.5±2.2 Hz to 26.6±3.6 Hz (W=171, z=3.058, p=0.002) as an effect of the simultaneous application of PHA-543613 and NS-1738. However, the simultaneous application of PHA-543613 and NS-1738 did not show higher relative effect than the sum of the relative firing rate changes during the individual applications of the two compounds (NMDA_PHA&NS-NMDA vs. (NMDA_PHA-NMDA)+(NMDA_NS-NMDA): 11.1±2.8 Hz vs. 12.9±4.0 Hz, W=107, z=0.483, p=0.629).

However, 5 out of the 6 neurons that responded with decreased NMDA-evoked firing rate to the single application of PHA-543613, showed an opposite response to the simultaneous application of PHA-543613 and NS-1738 by increasing their NMDA-evoked firing rate. We compared the sum of the effects of individual applications of PHA-543613 and NS-1738 with their co-application effect also in the restricted pool of those 6 neurons that showed decreased NMDA-evoked firing rate after PHA-543613 alone. The analysis revealed that the effect of the simultaneous application of PHA-543613 and NS-1738 increased the NMDA-evoked firing rate in a superadditive manner when the single application of PHA-543613 exerted an inhibitory effect (NMDA_PHA&NS-NMDA vs. (NMDA_PHA-NMDA)+(NMDA_NS-NMDA): −1.0±3.5 Hz vs. 16.6±5.9 Hz, W=21, z=2.201, p=0.028). This result suggests that the inhibitory action of PHA-543613 on the alpha7 nAChR may change to a facilitating effect when the PAM NS-1738 is also present.

## 4. Discussion

The present study aimed to address how modulation of alpha7 nAChRs by different pharmacological compounds influences firing activity and excitability of pyramidal neurons in the rat hippocampus. Our main finding was that the activation of alpha7 nAChRs by an orthogonal agonist or a PAM results in substantially different effects. Namely, firing rate increasing (facilitatory) and decreasing (inhibitory) effects were observed in almost equal number of neurons after the local application of the exogenous alpha7 nAChR agonist PHA-543613. The equal occurrence of facilitatory and inhibitory effects was typical for both spontaneous and NMDA-evoked firing activity. Earlier, Huang et al. (2010) investigated firing responses to alpha7 nAChR agonists in CA3 neurons, and they found solely facilitatory effects of the agonists when administered microiontophoretically. However, they also found 3 out of 10 neurons that decreased their firing rate after systemic administration of alpha7 nAChR agonist PSAB-OFP. Notably, substantial differences between the electrophysiological properties of CA3 and CA1 pyramidal cells might explain the dissimilar results (Rolls and Treves, 1997). Furthermore, we tested the effects of the alpha7 nAChR agonist PHA-543613 on a larger number of neurons which may have allowed us to detect such diverse effects with higher sensitivity. Nevertheless, the remarkable occurrence of the inhibitory effects of an alpha7 nAChR agonist is a novel and challenging finding. The most interesting finding is the inhibition of NMDA-evoked firing responses by the alpha7 nAChR agonist, an effect that has not been addressed in earlier *in vivo* reports, yet. Furthermore, the alpha7 nAChR agonist did not show overall synergistic interaction with the NMDA-induced firing rate increase; an interaction that has been earlier shown between NMDA and ACh and found to be dependent on alpha7 nAChRs (Bali *et al.*, 2017). These results suggest that direct targeting of alpha7 nAChRs with selective agonists does not perfectly mimic the alpha7 nAChR-dependent actions of the endogenous agonist ACh.

In contrast with PHA-543613, alpha7 nAChR PAMs predominantly increased the firing rate of the neurons and their responsiveness to NMDA and showed significantly higher increase of NMDA-evoked firing rate compared with PHA-543613. Furthermore, the PAM NS-1738 increased NMDA-responses in a superadditive manner, showing that the PAM facilitated the effects of endogenous ACh in the experimental arrangement applied here. These data fill a gap in the literature since there is only sparse earlier evidence on the *in vivo* electrophysiological effects of alpha7 PAMs, and no data is available on their specific effects on neuronal firing activity. However, alpha7 PAMs have been widely investigated in *in vitro* preparations that provide substantially different circumstances. In *in vitro* conditions, alpha7 PAMs do not evoke the opening of the channel pore, and no ionic current can be measured on alpha7 nAChRs during their sole application (Timmermann *et al.*, 2007; Sitzia *et al.*, 2011). However, both NS-1738 and PNU-120596 increases the peak current of ACh-evoked activation of alpha7 nAChRs. Furthermore, both compounds at least slightly modified the kinetics of receptor desensitization increasing the overall efficiency of receptor activation. In contrast with *in vitro* experiments, in the present study alpha7 PAMs exerted robust firing rate increasing effects alone without the application of any direct receptor agonist. These results suggest that there may be sufficient amount of endogenous ACh in the hippocampus of anesthetized rats to activate alpha7 nAChRs, and this effect can be further potentiated by the application of alpha7 PAMs. However, an earlier study found that in the presence of a PAM, alpha7 nAChRs on CA1 pyramidal cells can also be activated by the physiological level of endogenous alpha7 nAChR agonist choline (Kalappa *et al.*, 2010). The firing rate increasing effects of ACh on hippocampal pyramidal cells has been known for a long time, however, earlier results suggested that these effects are not mediated by nicotinic but only by the muscarinic ACh-receptors (Cole and Nicoll, 1984). In our previous report, we found that neither the ACh-evoked firing rate increase nor the NMDA-evoked firing rate increase was blocked by systemic administration of alpha7 nAChR antagonist MLA. On the other hand, synergistic effects of simultaneous cholinergic and glutamatergic activation was found to be dependent on the activation of alpha7 nAChRs (Bali *et al.*, 2017). Our results showing that alpha7 PAMs facilitated both spontaneous firing activity and responses to NMDA suggest that alpha7 nAChRs may essentially contribute to the cholinergic activation of CA1 pyramidal cells if the action of ACh on alpha7 nAChRs is amplified by a PAM.

Although earlier in vitro studies revealed that stimulation of alpha7 nAChR on stratum radiatum interneurons can shape pyramidal cell excitability through direct or indirect inhibition and disinhibition (Ji and Dani, 2000; Wanaverbecq et al., 2007), these indirect mechanisms are not likely to explain our present results because of two reasons. First, the recording electrode and iontophoretic drug delivery were located in the stratum pyramidale, where interneurons are less sensitive to nicotinic stimulation than in other layers of the hippocampus (McQuiston and Madison, 1999). Second, if distinct effects of the alpha7 nAChR agonist originated from indirect inhibitory/disinhibitory mechanisms through interneurons, then we would also expect to observe similar effects of the alpha7 nAChR PAMs on neuronal firing rate. On the contrary, the differential responses of neurons to alpha7 nAChR agonists and PAMs in the present study can be better explained by the rapid desensitization of the receptors following the opening of the channel by the binding of the direct agonist (Quick and Lester, 2002). Desensitization may explain that a substantial number of neurons responded with decreased firing rate and NMDA-evoked responses to PHA-543613 in contrast with alpha7 nAChR PAMs that exerted definite facilitatory effects in almost all neurons. In about half of all neurons, PHA-543613 presumably induced observable desensitization of the alpha7 nAChR receptors and prevented their activation by the endogenous agonist ACh, resulting in an antagonist-like inhibitory effect on the spontaneous and NMDA-evoked firing activity. The effects of PHA-543613 were comparable to those of alpha7 nAChR antagonist MLA as both compounds exerted increasing and decreasing effects on firing rate in similar number of neurons. On the contrary, while the distribution of the number of neurons with facilitatory or inhibitory effects of PHA-543613 was also equal on the NMDA-evoked firing activity, MLA predominantly and significantly decreased NMDA-evoked firing activity of the neurons. In summary, we found that the effects of alpha7 nAChR agonist PHA-543613 are different from the effects of both alpha7 nAChR antagonist MLA and PAMs NS-1738 and PNU-120596 on the level of neuronal firing activity.

Furthermore, co-application of alpha7 nAChR agonist PHA-543613 and PAM NS-1738 resulted in additive effects on both spontaneous and NMDA-evoked firing activity. Moreover, NS-1738 counteracted the inhibitory effect of PHA-543613 in most neurons that responded with decreased NMDA-responses to PHA-543613 application resulting in a superadditive increase of the NMDA-evoked firing rate during co-application. This result is also in line with the hypothesis that the inhibitory effects of PHA-543613 may have been originated in the desensitization of the alpha7 nAChRs. Although usually categorized as a Type I PAM, NS-1738 does not only increase the agonist-evoked current on alpha7 nAChRs, but it also has a marginal blocking effect on the desensitization of the receptors according to *in vitro* experiments (Timmermann *et al.*, 2007). Thus, in the presence of NS-1738, PHA-543613 may have caused weaker desensitization of alpha7 nAChRs and the activation of the receptors may have become predominant in the effect of PHA-543613 on the NMDA-evoked firing rate.

There is a large body of evidence for the cognitive enhancing effects of both alpha7 nAChR agonists and PAMs (for review, see Pandya and Yakel (2013), and Wallace and Porter (2011)). Specifically, PHA-543613, NS-1738 and PNU-120596 were also found effective in improving cognitive performance in different behavioral paradigms in different animal models (Timmermann *et al.*, 2007; Bali *et al.*, 2015; Nikiforuk *et al.*, 2015; Sadigh-Eteghad *et al.*, 2015; Nikiforuk, Kos, *et al.*, 2016). At the moment, it is still not possible to decide whether the desensitization of alpha7 nAChRs decreases the efficacy of agonist compounds in cognitive enhancement. However, it is suggested that tolerance may evolve against alpha7 nAChR agonists because of the desensitization of the receptors, therefore, alpha7 nAChR PAMs are considered to provide better therapeutic perspectives for cognitive impairment (Corradi and Bouzat, 2016). According to our present results, PAMs more potently increased neuronal excitability in the hippocampal CA1 area than alpha7 nAChR agonists, which may imply higher acute cognitive enhancer potential. However, some reports suggest that the inhibition of receptor desensitization by Type II PAMs of alpha7 nAChRs induce neuronal cell death by overly increasing intracellular Ca2+ levels (Guerra-Álvarez *et al.*, 2015). Thus, it is still questioned whether or not the blocking of receptor desensitization is a beneficial action of potential medications for neurocognitive disorders (Williams *et al.*, 2011). In this regard, the inhibitory effects of PHA-543613 on NMDA-evoked firing rate in our present experiment may be considered as a protective effect against glutamatergic overstimulation of pyramidal cells. On the other hand, NS-1738 and PNU-120596 did not show such inhibitory effects, suggesting that alpha7 nAChR PAMs increase neuronal excitability without limitation and protection from overstimulation. Thus, desensitizing alpha7 nAChR agonists may be considered as neuroprotective compounds, while blocking desensitization by PAMs may result in certain neurotoxic effects in a long term application. However, this presumption needs further assessment with different neurophysiological and neuropharmacological methods on the cellular and behavioral levels.

Furthermore, we observed a positive interaction between the effects of PHA-543613 and NS-1738 on the NMDA-evoked firing rate. There is only sparse evidence in the literature about the synergistic interaction of alpha7 nAChR agonists and PAMs in functional tests. Freitas et al. (2013) found that an alpha7 nAChR agonist (choline or PHA-543613) and a PAM (PNU-120596) synergistically reduced formalin-induced pain following their co-administration in mice. Furthermore, alpha7 nAChR PAM PNU-120596 reportedly facilitated the cognitive enhancer effect of AChE inhibitor donepezil in rodents and non-human primates (Callahan *et al.*, 2012). This finding is in line with our present results that alpha7 nAChR PAMs increase firing rate when administered alone, and positively interact with the alpha7 nAChR agonist during their co-application. Taken together, it is reasonable to hypothesize that increased levels of ACh as an effect of AChE inhibitors in combination with the administration of alpha7 PAM compounds may also additively or synergistically increase the firing rate and NMDA-evoked responses of the neurons providing a basis for synergistic behavioral effects.

## 5. Acknowledgements

The authors would like to thank Balázs Knakker and Evelin Kiefer for reviewing the manuscript and giving valuable suggestions. The authors also thank Mrs. Jánosné Antal for her assistance in animal care.

## Abbreviations

ACh: acetylcholine
AChE: acetylcholinesterase
nAChR: nicotinic acetylcholine receptor
MLA: methyllycaconitine
NMDA: N-methyl-D-aspartate
NMDAR: N-methyl-D-aspartate receptor
PAM: positive allosteric modulator

